# Bacteriophages dynamically modulate the gut microbiota and metabolome

**DOI:** 10.1101/454579

**Authors:** Bryan B. Hsu, Travis E. Gibson, Vladimir Yeliseyev, Qing Liu, Lynn Bry, Pamela A. Silver, Georg K. Gerber

## Abstract

The human gut microbiome is comprised of densely colonizing micro-organisms in dynamic interaction with each other and the host. While the bacterial component of the microbiome is under intense investigation, far less is known about how bacteriophages impact bacterial communities in the gut. We investigated the dynamic effects of phages on a model microbiome using gnotobiotic mice colonized by commensal bacteria that colonize the human infant gut, and found that phage predation not only directly impacts susceptible bacteria but also leads to cascading effects on other bacterial species *via* inter-bacterial interactions. Using metabolomic profiling, we also found that the shifts in the microbiome caused by phage predation have a direct consequence on the gut metabolome. Our work provides insight into the ecological importance of phages as modulators of bacterial colonization, and additionally suggests the potential impact of gut phages on the host with implications for the use of phages as therapeutic tools to rationally and precisely modulate the microbiome.

## Introduction

Our bodies contain roughly as many bacterial cells as our own, with the greatest density in the colon at 10^14^ cells (Sender et al., 2016). This microbiome benefits human health *via* mechanisms including metabolic cross-feeding and immune system priming (Brestoff and Artis, 2013). Conversely, an imbalanced or depleted microbiome can be deleterious. Diseases associated with an abnormal microbiome include poor nutrient utilization (Smith et al., 2013; Turnbaugh et al., 2006), gastrointestinal disease (Chang et al., 2008; Frank et al., 2007), as well as diseases of the liver (Le Roy et al., 2013), heart (Zhu et al., 2016), and brain (Hsiao et al., 2013). As we gain greater understanding of the role of the gut microbiome in human health and disease, the leading question becomes: what factors in the gut influence the microbes that influence us, and how can we leverage this knowledge toward treatments that manipulate the microbiome?

Current approaches to modulating the gut microbiome include dietary changes and antibiotics, but these modalities cause broad and potentially long-lasting perturbations. For infants, whose gut microbiomes are in a nascent state, the consequences of perturbations are of particular concern as it can lead to depleted bacterial diversity and instability in the short term (Yassour et al., 2016) and increased incidences of allergic disease (Bisgaard et al., 2011), overweightness (Ajslev et al., 2011) and asthma (Abrahamsson et al., 2014) in the long term. From both the basic scientific and therapeutic perspectives, strategies are thus needed to more precisely and rationally modulate gut bacteria within a complex community (Schmidt et al., 2018). A promising approach toward this goal is to study the ecological antagonists of bacteria in the gut, bacteriophages (phages). Analogous to studying natural products or their derivatives for therapeutic purposes (Dias et al., 2012), studying the role of bacteriophages in the gut may illuminate new approaches for: (1) deliberately and rationally perturbing specific bacteria, (2) elucidating causality as mediated by inter-bacterial and bacterial-host interactions, and (3) ultimately designing precise and predictable approaches to remodel the gut microbiota for therapeutic purposes.

Phages are prokaryotic viruses that are among the most abundant microbes in the gut, but also among the least understood (Keen and Dantas, 2018; Mirzaei and Maurice, 2017). These viruses generally propagate *via* lytic or lysogenic infection of bacteria, often with species-level specificity. While their metagenomic composition has been associated with diseases such as inflammatory bowel disease (Norman et al., 2015) and malnutrition (Reyes et al., 2015), little is known about the actual behavior of phages in the gut (Reyes et al., 2013). Analogous to the importance of understanding apex predators in natural environments (Estes et al., 2011), elucidating the predation behavior of phages has the potential to provide similar insights in the gastrointestinal ecosystem. However, tracking the cause-effect relationship of phage predation on naturally-occurring microbiomes and consequently on host and microbial metabolism is extremely challenging (Fischbach, 2018; Schmidt et al., 2018). Gnotobiotic mice, colonized with a limited and known but still complex collection of bacteria, present an attractive model system for comprehensively characterizing the behavior of phages in the gut environment.

In this work, we administered lytic phages to gnotobiotic mice colonized with a defined set of human commensal bacteria and longitudinally-tracked the response of each microbe using high-throughput sequencing and targeted molecular methods. We found that phages cause targeted knockdown of susceptible species in the gut, and further modulate non-targeted bacteria through inter-bacterial interactions, resulting in blooms and attrition of previously repressed and promoted species. By comparing the colonization profile of a full bacterial consortia to those omitting phage-targeted species, we delineate the causal effects of simultaneous phage predation. Using broad metabolic profiling, we demonstrate that the compositional shifts in bacteria caused by phage predation also modulate the gut metabolome. These findings have implications for effects on the host, and the potential use of phages for therapeutic purposes.

## Results

### Phages are specific for individual species among representative human gut commensals

Because the infant gut microbiome is especially vulnerable to perturbations with long-term consequences, we constructed a model microbiota comprised of human facultative- and obligate-anaerobic commensal bacterial species known to colonize the infant human gut (Blanton et al., 2016; Gibson et al., 2016; Smith et al., 2013; Subramanian et al., 2014), and that also can stably co-colonize germ-free mice (Bucci et al., 2016). The ten selected species represent the major phyla in the human gut microbiome (The Human Microbiome Project Consortium, 2012), namely Firmicutes (*Clostridium sporogenes*, *Enterococcus faecalis*), Bacteroidetes (*Bacteroides fragilis*, *Bacteroides ovatus*, *Bacteroides vulgatus*, and *Parabacteroides distasonis*), Proteobacteria (*Klebsiella oxytoca*, *Proteus mirabilis*, and *Escherichia coli* Nissle 1917), and Verrucomicrobia (*Akkermansia muciniphila*). Each of the species selected are also readily available from strain collections, culturable *in vitro*, and genetically characterized. For four of these species, we selected lytic phages because of their availability in microbiological repositories and past description in literature, namely *E. coli* (T4 phage (Miller et al., 2003)), *C. sporogenes* (F1 phage (Betz and Anderson, 1964)), *B. fragilis* (B40-8 phage (Tartera and Jofre, 1987)), and *E. faecalis* (VD13 phage (Ackermann et al., 1975)).

We first confirmed the specificity of our phages by testing their ability to lyse a panel of our candidate, and putatively non-susceptible, human gut commensal bacteria. Using a spot assay (Kutter, 2009), we tested the susceptibility of each bacteria to 5 µL of each lytic phage (∼10^9^ pfu/mL). After incubation at 37°C, aerobically or anaerobically depending on bacterial culture conditions, we found that T4, F1, B40-8, and VD13 phages lysed only their susceptible bacteria and had no apparent impact on the other commensal bacteria (Figure S1, Table S1).

### Phages knockdown and co-exist with susceptible bacteria in the mammalian gut

With little known about the dynamics of phage predation in the mammalian gut, we first aimed to characterize the kinetic behavior of both phages and their targeted bacteria in the context of a larger bacterial community. As shown in Figure 1A, we colonized germfree mice with our defined bacterial consortia of ten species and then introduced phages targeting a subset of these species in the following sequence: T4 and F1 phages targeting *E. coli* and *C. sporogenes*, respectively; an equilibration period of two weeks; B40-8 and VD13 phages targeting *B. fragilis* and *E. faecalis*, respectively; and a final equilibration period of two weeks. Phages were administered in pairs to probe whether multiple simultaneous perturbations could potentially have synergistic or nullifying effects. Each set contained phage targeting a facultative- and obligate-anaerobe to reduce the potential bias of one phage set over the other. Stool was regularly collected, with greater frequency around perturbations to capture the information-rich dynamical changes (Gerber et al., 2012), and analyzed for microbial composition using a combination of quantitative PCR and high-throughput sequencing of 16s rRNA amplicons.

**Figure 1.**
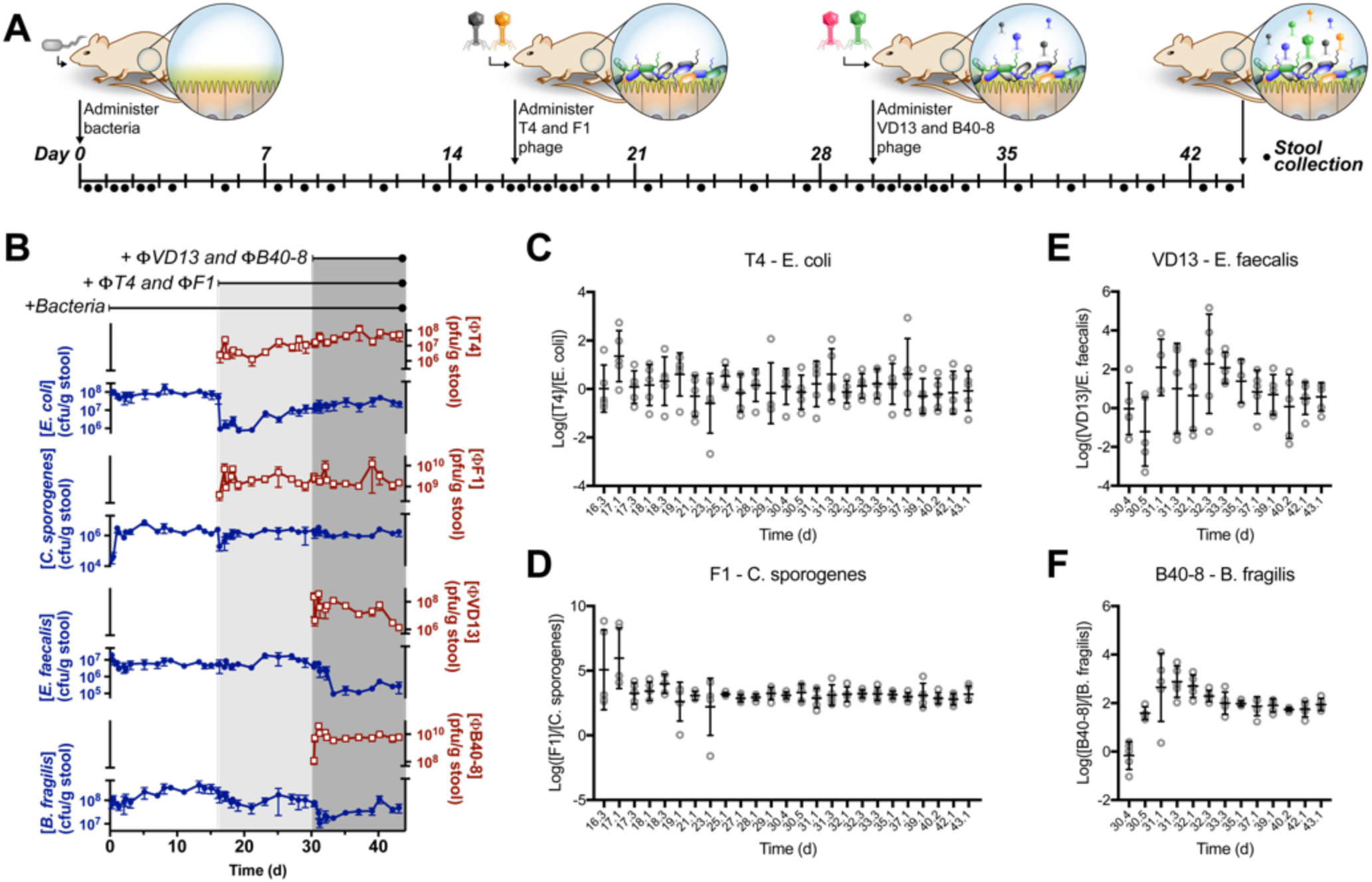
Longitudinal characterization of phage behavior in the mammalian gut. (A) Individually housed germfree C57Bl/6 mice (n=5) were orally gavaged with a bacterial mixture containing ∼10^6^-10^7^ cfu/strain. On Day 16.1 and 30.1, mice were gavaged with 1 M of NaHCO_3_ to neutralize gastric acid, followed by ∼10^6^ pfu/phage. Fecal samples were collected periodically throughout the experiment, with increased frequency around instances of perturbation. (B) Concentrations of each phage and targeted bacteria shown as (pfu/g stool) and (cfu/g stool), respectively. (C) Relative ratios (log_10_) of phage per bacteria derived from data in Figure 1B are shown for T4 phage and *E. coli*, (D) F1 phage and *C. sporogenes*, (E) VD13 phage and *E. faecalis*, and (F) B40-8 phage and *B. fragilis* in stool.

Our analysis revealed that both phages and their targeted commensal bacteria persist in the gut. After administration, each phage was detectable within ∼4-6 h, and continued to be detectable for the duration of the experiment (Figure 1B). As non-replicating phage is generally shed completely within the first couple of days (Weiss et al., 2009), the continued presence of each phage in the gut indicates that the rate of phage production is sufficient to overcome the decay rate from inactivation and intestinal emptying.

We further found that phage and bacteria demonstrate a dynamical interaction during the initiation of predation which may provide a window into the *in situ* properties of phage propagation. The relative number of phage to bacteria (*i.e.*, multiplicity of infection) can determine the kinetics during the early stages of phage infection. The dynamics of B40-8—*B. fragilis* reveals an initially low fraction of phage per bacteria that peaks and gradually stabilizes to an intermediate level (Figure 1F). We see similar results for VD13—*E. faecalis* (Figure 1E), although there is considerable mouse-to-mouse variability in this case. In contrast, the dynamics of T4—*E. coli* appear to immediately reach steady state within the first sampling (Figure 1C). Less is clear for F1—*C. sporogenes* as a singular peak in the trajectories for four of five mice appears within the first day after phage administration (+0.2d and +1d) (Figure 1D). These differential dynamics between each phage-bacteria interaction could be indicative of unique properties of phage propagation or the accessibility of bacteria to infection in their local environments.

### Phages induce cascading effects in species in the microbiota not directly targeted

As shown in Figure 2A, the introduction of phages caused quantitative shifts in bacterial colonization density beyond the target species. After the first set of phage, administered on Day 16.1, *A muciniphila* blooms to be one of the more abundant species coinciding with a depletion of *B. fragilis.* The lower-abundance species are also modulated as *B. vulgatus, P. mirabilis*, and *P. distasonis* are also enriched after this modulation. Less obvious are the effects due to the second set of phage administered on Day 30.1, even when examining the trajectories of each species separately (Figure S2). By examining the relative changes in bacterial colonization resulting from phage administration as shown in Figure 2B, we found that the knockdown of *C. sporogenes* and *E. coli* during the first set of phage administrations results in a rapid and substantial expansion of *B. vulgatus*, *P. mirabilis*, and *A. muciniphila*, followed by a gradual expansion of *P. distasonis* and *B. ovatus*, and a gradual reduction of *B. fragilis*. The impact of the second set of phages on the surrounding microbiota was less pronounced, with minimal expansion of *E. coli* and *C. sporogenes* and fluctuating behavior from *B. ovatus*, *B. vulgatus*, and *K. oxytoca*. Trajectories of other species were near baseline (Figure S3).

**Figure 2.**
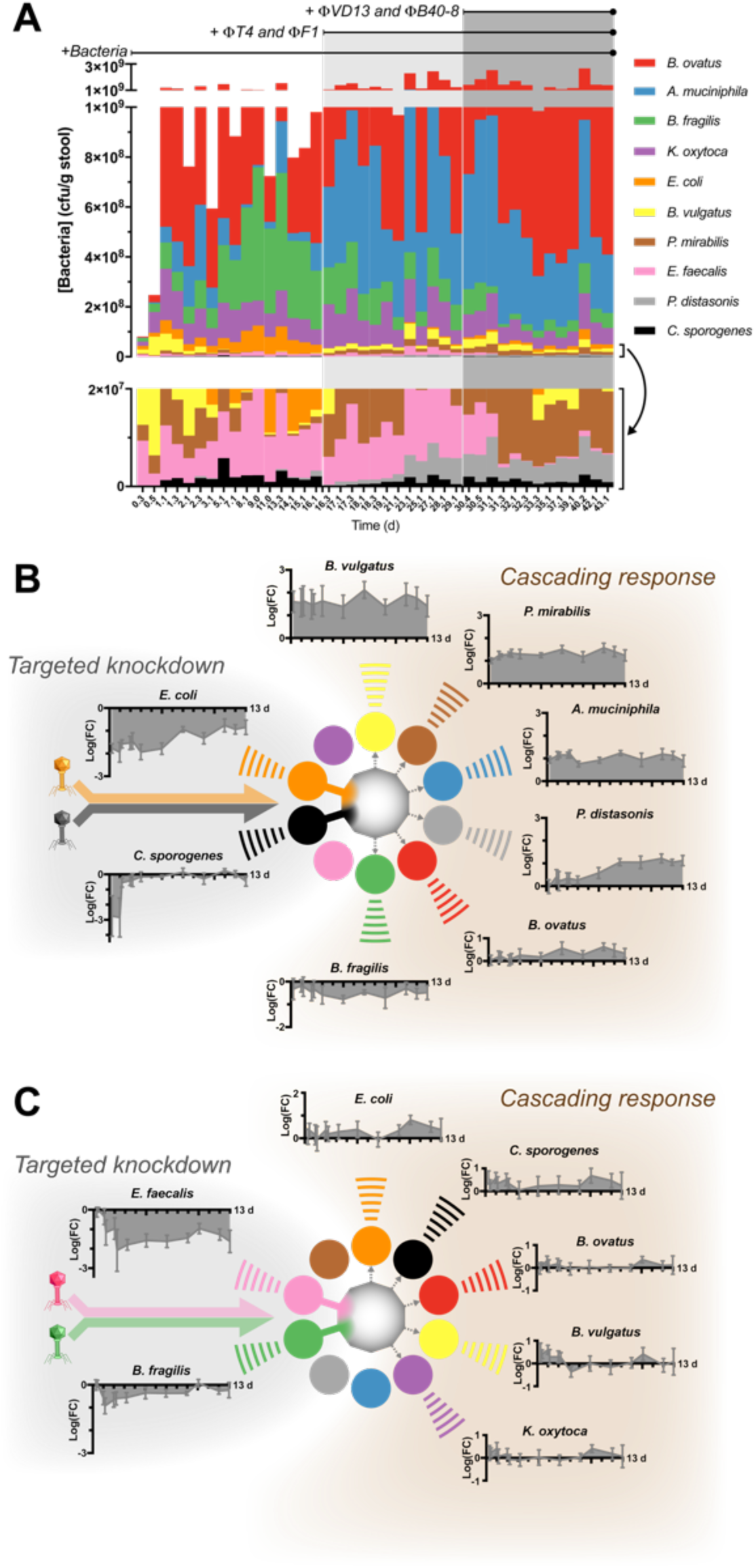
Effect of phage on the commensal gut microbiota. (A) Fecal abundances of bacteria with the lower panel as a zoomed-in version of the top full-scale panel. (B) Relative fold-changes (log_10_) in each bacterial species, derived from data shown in Figure 2A, resulting from introduction of the first set of phage, and (C) second set of phage.

Despite the modulation of the background flora, phage predation did not induce changes in species richness. As shown compiled in Figure 2A and individually in Figure S2, none of the bacterial species are completely eliminated after introduction of phage. In fact, the overall bacterial load does not change markedly (Figure S4). It has been suggested that the presence of predators can engender greater stability to the ecosystem (Allesina and Tang, 2012). Calculations of Bray-Curtis similarity in our defined microbiota reveals an increased similarity across mice (Figure S5A) and between adjacent time points with each successive introduction of phage (Figure S5B). Increased similarity across mice and between time points is indirect evidence of a more stable system that reduces deviations from steady state, suggesting that phage predation may benefit bacterial communities in the gut.

### Knockout experiments of bacteria targeted by phages delineate causal effects of bacterial interactions

Comparing the colonization of mice with and without each phage-targeted species revealed its quantifiable impact on the surrounding microbiota. As conceptually outlined in Figure 3A, comparing bacterial colonization in the presence of *Species 1* (full consortia) to its absence (knockout) yields insights into its facilitative and inhibitory roles on the surrounding microbiota. Consequently, attrition of *Species 2* colonization in the knockout is indicative of lost promotion from *Species 1* and conversely the bloom of *Species 3* in the knockout represents lost repression from *Species 1*. As phage predation equates to bacterial knockdown, its impact will approach that of the knockout (Figure 3A). To this end, we quantified the colonization densities of knockout consortia (Figure 3B) by inoculating four sets of germfree mice with a bacterial consortia comprising nine species of the original ten-member consortia described earlier, omitting either *E. coli, C. sporogenes, B. fragilis*, or *E. faecalis* in turn. As shown in Figure 3C, the colonization profile for each cohort of mice revealed marked compositional differences compared to other dropouts and with the full consortia (corresponding data from Figure 2A) despite similar overall mean bacterial densities at steady state: full consortia = 1.1 × 10^9^ cfu/g stool, *E. coli* dropout = 6.9 × 10^8^, *C. sporogenes* dropout = 1.1 × 10^9^, *B. fragilis* dropout = 9.1 × 10^8^, and *E. faecalis* dropout = 7.3 × 10^8^. We calculated the influence of each omitted species by comparing the colonization density of each species in its dropout to the full consortia at steady state (Day 16). As shown in Figure 3D, each omitted species has a distinct pattern of influence on the defined consortia, which indicates that the knockdown of each species by phage should produce a phenotypically unique result.

**Figure 3.**
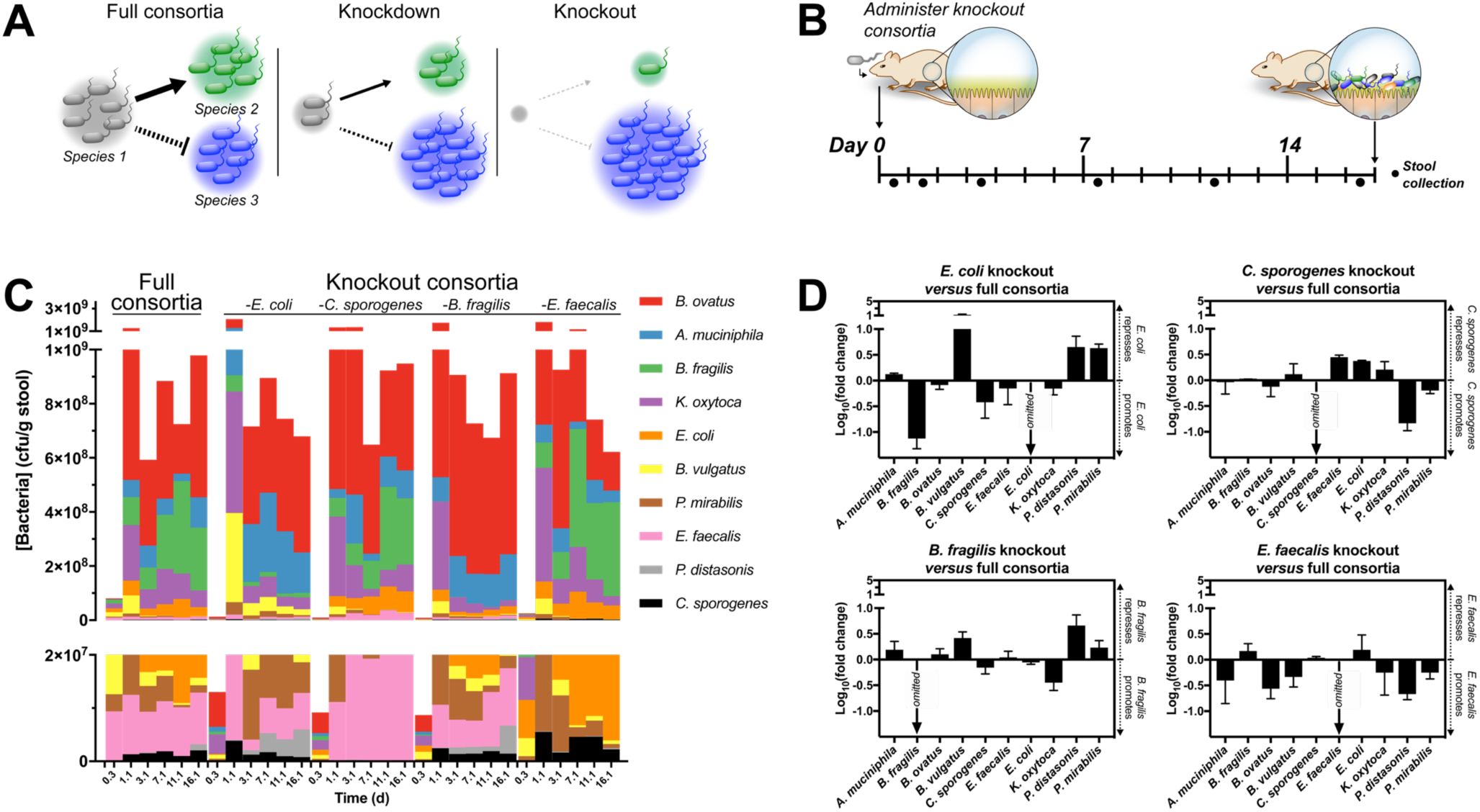
Impact of bacterial knockouts on colonization of the consortia. (A) Schematic representation of how colonization densities of the surrounding microbiota respond to the omission of individual species (*Species 1*) in bacterial knockout consortia. (B) Four sets of five individually housed germfree C57Bl/6 mice were orally gavaged with a bacterial mixture (∼10^6^-10^7^ cfu/strain) containing nine of the original ten-member consortia, omitting each of *E. coli, C. sporogenes, B. frailis*, and *E. faecalis* in turn. (C) Colonization densities of either a complete (“full”) or knockout consortia. (D) Relative fold change (log_10_) in colonization by each species in dropout consortia normalized to full consortia.

Using the magnitude of relative changes in colonization with and without phage-targeted species (Figure 3D), we hypothesize a bacterial interaction network. To represent the effects of our phage administration, we conflated the effects of the first set of phage (T4 and F1 phages targeting *E. coli* and *C. sporogenes*, respectively) and second set of phage (B40-8 and VD-13 phages targeting *B. fragilis* and *E. faecalis*, respectively), as shown in Figure 4A and 4B, respectively. Because bacterial omission provides the maximal effect possible, the magnitude of inhibitory or facilitative interactions within this interaction network provides an upper-limit to the impact of phage predation, which knocks down but does not eliminate the target bacteria.

**Figure 4.**
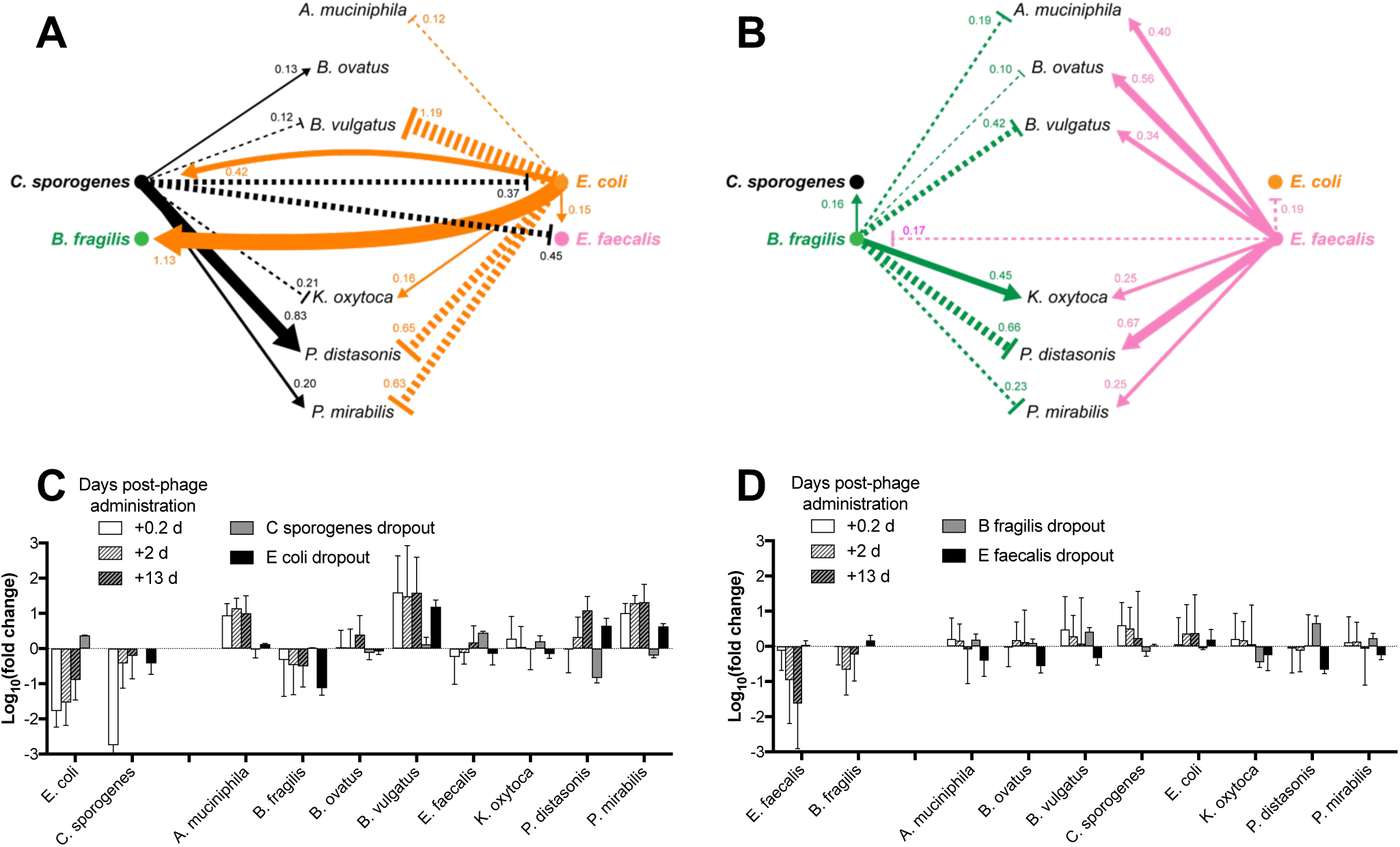
Bacterial interaction network in the gut microbiota. Hypothesized causal interaction network derived from differences in colonization between the full ten member consortia and dropouts of phage-targeted bacteria at steady state (Day 16). Linewidths correspond to the knockout-induced change in colonization densities (*i.e.*, log_10_ (fold change). (A) Network representing the combined effects of *E. coli* and *C. sporogenes* or (B) *E. faecalis* and *B. fragilis*. (C) Log_10_(fold change) in concentration of each species from mice colonized with the full consortia (data from Figure 2) at select time-points for comparison.

In some cases, phage-directed knockdown can have a subtractive effect that approaches that of bacterial omission, as exemplified by T4 phage, which substantially reduces the population of *E. coli* immediately after administration. As indicated by our interaction network (Figure 4A), *E. coli* strongly promotes *B. fragilis* and strongly represses *B. vulgatus* through bacterial interactions and thus its knockdown by the first set of phages leads to a contraction in *B. fragilis* and a bloom of *B. vulgatus.* The magnitude of these effects approaches the levels observed in the dropout studies (Figure 4C), likely due to *E. coli* being the primary influencer of these species.

Causal effects become less clear when two counteracting effects influence a single species, but can still be rationalized using the interaction network. For example, after the administration of our second set of phages that targeted *E. faecalis* and *B. fragilis*, we found a relatively minimal impact on the colonization of the surrounding microbiota (Figure 2C). However, our interaction network for the second set of phage targeted species (Figure 4B) revealed that *A. muciniphila, B. ovatus, B. vulgatus, P. distasonis*, and *P. mirabilis* are all repressed by *B. fragilis* and promoted by *E. faecalis*. Thus, the simultaneous knockdown of these two phage-targeted species during the second set of phage administered resulted in counteracting losses in repression and promotion, ultimately nullifying their individual effects on the community and leading to a negligible impact on over all bacterial colonization.

We also observed interesting temporal dynamic effects on the microbes due to phage-predation, which can be explained by bacterial interactions. For example, we observed that *P. distasonis* had a delayed bloom that began approximately 3 days after phage administration (Figure 2B), despite other species demonstrating an immediate response. Bacterial interactions as depicted in Figure 4A suggest a mechanism for this behavior. Soon after introduction of the first set of phage, *C. sporogenes* and *E. coli* are knocked down (Figure 1B), which results in a loss of their respective promotion and repression of *P. distasonis*. With the sustained knockdown of *E. coli*, *P. distasonis* experiences a similarly durable bloom due to the de-repression. However, *C. sporogenes* only experiences an initial transient knockdown and its recovery after the first few days coincides with the rescue of its promotion of *P. distasonis*, thus, explaining a second bloom of *P. distasonis* beginning after day 3. This effect is similarly observed for *P. mirabilis*, though to a lesser degree because of the weaker promotion by *C. sporogenes*.

Our results also suggest some deeper cascading effects not captured in our interaction network derived from the dropout experiments. As shown in Figure 4C, knockdown of *E. coli* and *C. sporogenes* by the first set of phage leads to an enrichment of *A. muciniphila* substantially beyond what is described by the dropout experiments. As our study examined the causal effects of four of the ten members of the consortia, it is likely that *A. muciniphila* experiences additional influence from other members of the microbiota (*e.g., B. vulgatus, B. ovatus, P. distasonis*, and/or *P. mirabilis*).

### Bacterial modulation induced by phages impacts the gut metabolome

We sought to characterize the functional effects of phage-predation on the microbiome as reflected in changes in the gut metabolome. Overall, our expectation was that levels of most metabolites would be buffered against fluctuation due to metabolic redundancy across our defined consortia, but compounds associated with microbial pathways unique to particular species in our consortium would be sensitive to perturbations in the bacterial composition. Using untargeted metabolomics, we surveyed the fecal metabolites during various stages of colonization in mice, namely sterile (germfree), after stable bacterial colonization, after introduction of *E. coli* and *C. sporogenes* phages, and after introduction of *E. faecalis* and *B. fragilis* phages (Figure 5A).

**Figure 5.**
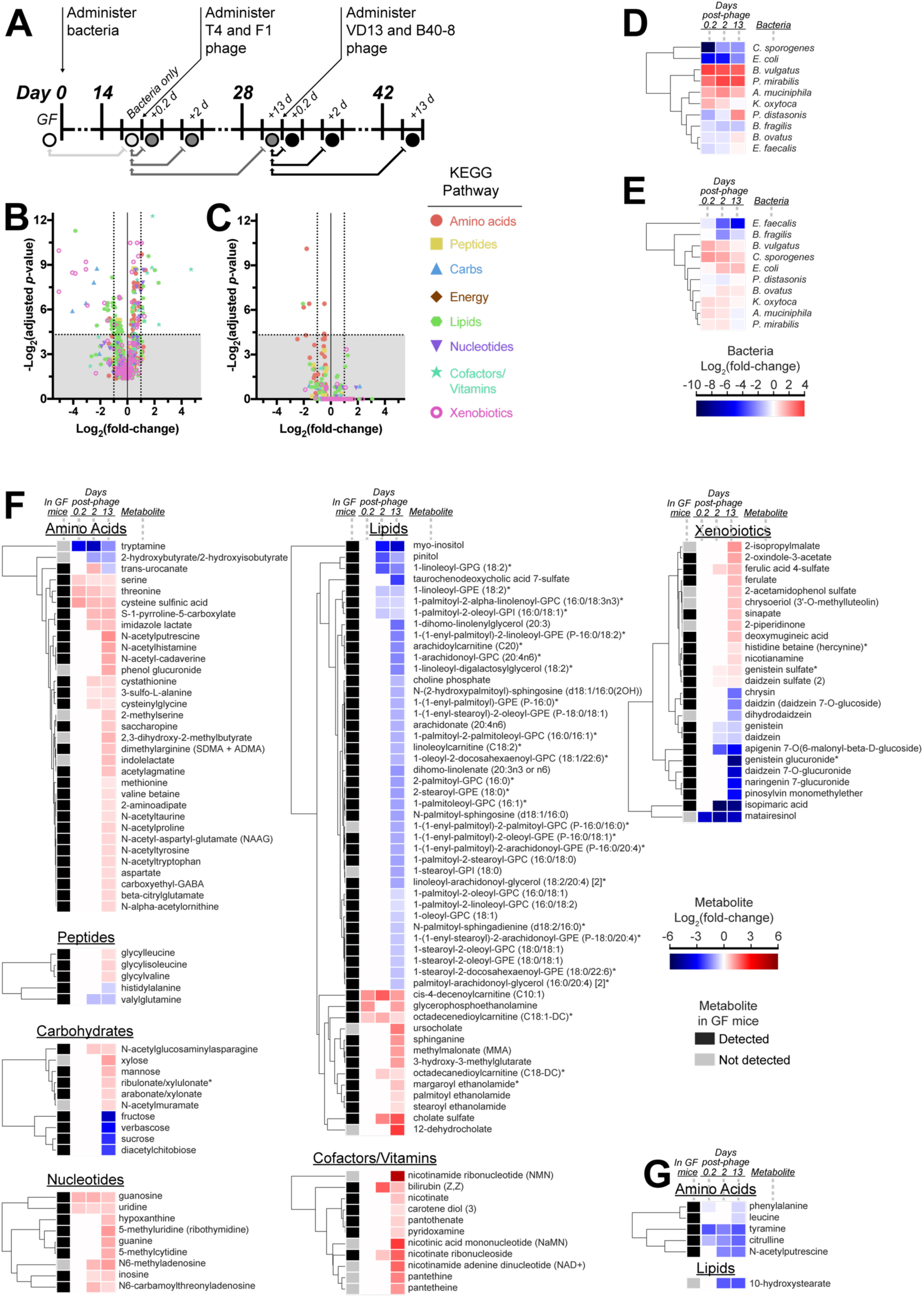
Analysis of the fecal metabolome. (A) Fecal samples were collected from GF mice colonized by the defined bacterial consortia, then treated with the first set of phages (T4 and F1) and second set of phages (VD13 and B40-8). Relative concentrations of metabolites were normalized to values from samples immediately prior to phage administration. (B) Volcano plots showing increasing significance (y-axis) versus fold change (x-axis) of each metabolite 13 days after administration of the first set of phage and (C) each metabolite 13 days after administration of the second set of phage. Points above the horizontal dashed line indicate significant changes with adjusted *p*-values < 0.05. (D) Hierarchical clustering of each bacterial species after introduction of the first set of phage and (E) the second set of phage. (F) Hierarchical clustering of significantly changing (adjusted *p*-values < 0.05) metabolites after introduction of the first set of phage and (G) the second set of phage. Cutoff for presence of each metabolite in GF mice was detection in at least 4 of 5 mice.

Overall, phage-directed remodeling of the gut microbiota had a relatively narrow impact on the metabolome. Administration of the first set of phages resulted in statistically significant changes in 17% of examined compounds, with metabolites representing nearly all the KEGG pathways (*e.g.*, amino acids, peptides, carbohydrates, lipids, nucleotides, cofactors/vitamins and xenobiotics) (Figure 5B and Table S2). We also found that the second set of phage had a comparatively limited impact, as only 0.7% of metabolites were significantly affected (Figure 5C and Table S2), which coincided with a relatively limited change in the microbiota that was dominated by a decrease in *E. faecalis* 13 days post-phage (Figure 5E). By comparison, introduction of bacteria to germfree mice resulted in broad shifts in the fecal metabolome, enriching 60% and reducing 15% of metabolites across all KEGG pathways (Figure S6 and Table S2). Taken together, these observations suggest that the breadth of metabolomic impact mirrors the extent of compositional shift in the gut microbiota.

### Phages can modulate neurotransmitter metabolites uniquely associated with specific bacteria

We observed that in some cases the specificity of phage-predation allows for the targeting of bacterial species and consequently the knockdown of uniquely-associated metabolic products. Tryptamine is a neurotransmitter commonly of plant origin but is also produced via tryptophan decarboxylation by a small number of commensal gut bacteria. While this gene can be found in ∼10% of human gut bacterial metagenomes, it has so far been only identified in two genetically-characterized species, *R. gnavus* and *C. sporogenes* (Williams et al., 2014), the latter of which is a member of our defined consortia. BLAST search of tryptophan decarboxylase amino acid sequences from *R. gnavus* (rumgna_01526) and *C. sporogenes* (clospo_02083) against the other members of our consortia showed poor protein homology (top hit of 31% identity, Table S3A), consistent with its unique association to *C. sporogenes*. During treatment with the first set of phage, we detected a 10-, 17-, and 2-fold reduction in tryptamine (0.2, 2, and 13 days, respectively) as shown in the amino acid pathway depicted in Figure 5F. This corresponds with an 840-, 4-, and 4-fold reduction in *C. sporogenes*, respectively (Figure 5D).

As another example, the neurotransmitter tyramine is produced via tyrosine decarboxylation by lactic acid bacteria including *E. faecalis* (Connil et al., 2002), the sole lactic acid bacteria of our consortia. We found no associations of tyrosine decarboxylase with other consortia members in the literature nor any significant protein homology to the *E. faecalis* protein (tyrDC) by BLAST (top hit of 28% identity, Table S3B), consistent with tyrosine decarboxylation function solely associated with *E. faecalis*. Administration of our second set of phage caused a 4-, 2.7-, and 4-fold decrease in tyramine (0.2, 2, and 13 days, respectively) as shown in the amino acid pathway of Figure 5G. This corresponds with a 1.3-, 9-, and 42-fold reduction in *E. faecalis*, respectively (Figure 5E). Due to the limited catalog of experimentally verified microbial metabolites (Dorrestein et al., 2014), it is difficult to broadly associate specific metabolites to individual species within our consortia. However, the unique associations of tryptamine and tyramine to *C. sporogenes* and *E. faecalis*, respectively, suggest clear causal links between phage, bacteria, and metabolite.

### Phages can modulate metabolites with known host effects associated with multiple bacterial species

Compounds more broadly associated with microbial metabolism were also significantly impacted by phage-directed shifts in the microbiota. For example, the first set of phages also increased fecal concentrations of two amino acids, serine and threonine, which are highly-represented amino acids of *O*-glycosylated intestinal mucin (Derrien et al., 2010), consistent with our observed enrichment of the mucin-degrading commensals *A. muciniphila* and *B. vulgatus*.

We also found significant changes in bile salts due to the first set of phage. Tauro- and glyco-conjugated primary bile salts produced by the host undergo microbial transformations including amino acid deconjugation by bile salt hydrolases (BSH) and dehydrogenation by hydroxysteroid dehydrogenases (HSDH). We found that modulation of the gut consortia by the first pair of phages increased the deconjugated bile salt, cholate sulfate, and decreased the conjugated bile salt, taurochenodeoxycholic acid 7-sulfate (Figure 5F). This suggests an increased activity of BSH which we found prevalently associated with our consortia (*B. fragilis, B. ovatus, B. vulgatus, C. sporogenes, E. faecalis, E. coli, P. distasonis*, and *P. mirabilis*) as described in the MetaCyc database (Caspi et al., 2016). We also detected increases in two deconjugated, secondary bile salts that were not detected in germfree mice and thus microbially derived: 12-dehydrocholate and ursocholate. The former is produced by 12a-HSDH activity while the latter is produced by sequential 7a-HDSH and 7b-HDSH activity. Counterintuitively, each enzyme is associated with *B. fragilis*, *C. sporogenes*, and *E. coli* (Caspi et al., 2016), three species that correspondingly decrease after the first set of phage. Other factors are likely involved, including changes in bile salt absorption by the host and the capability of other consortia members for bile salt metabolism that have yet to be experimentally characterized.

## Discussion

Our results demonstrate that lytic phages not only knockdown their bacterial targets, but also affect non-susceptible species within a community of commensal bacteria colonizing the gut through cascading effects. We verified phage-bacteria interactions *in vitro* and then longitudinally-followed their impact on a defined gut microbiota in a gnotobiotic mouse model. Using knockout experiments in which mice were colonized with all but one member of our defined microbiota, we mapped inter-bacterial interactions and elucidated the causal effects of phage perturbation in our defined microbiota. Our results depict a highly interactive and dynamic community where lytic phage coexist and knockdown targeted bacteria, with an effect that propagates through the other members of the microbiota to ultimately modulate the gut metabolome.

Although phage-predation has classically been viewed through the lens of species- or strain-specific impact on bacteria, our results highlight the importance of considering inter-bacterial interactions within bacterial communities, as perturbations of individual species can have cascading effects. Our findings are consistent with the emerging understanding of how the gastrointestinal environment (*e.g.*, dense colonization, niche competition and nutrient limitations) promotes intense competition and cooperation among species (Hibbing et al., 2010). With *in vitro* verification of phage-bacteria interactions, we could analytically separate the effects of phage directed knockdown from the subsequent modulation of the microbiota through inter-bacterial interactions. As highlighted in Figure 2, it is clear that targeted modulation of bacterial species has a subsequent effect across co-colonizing species in the gut.

While clearly important, inter-bacterial interactions are challenging to experimentally identify and confirm *in vivo* given the limited tools currently available (Schmidt et al., 2018). By analogy, in molecular biology, a general strategy to confirm the putative role of a gene is to verify loss-of-function using a genetic knockout and then verify gain-of-function by reintroducing the gene to the knockout. Our results suggest that phages can provide similar information for the microbiome, although in a graded rather than absolute manner. For example, sustained knockdown of *E. coli* using T4 phage results in sustained modulation of the surrounding gut bacteria, effects that approach those observed when *E. coli* was completely omitted from colonization in the dropout experiments. Similarly, the dynamics of predation by F1 phage, where *C. sporogenes* is initially knocked down but rapidly recovers within days, provides “knockdown” and “overgrowth” information which we found had a functional effect represented by the delayed bloom in *P. distasonis*. By extension, the rational deployment of multiple phages could be designed to selectively modulate certain species while minimizing the cascading influence on the surrounding microbiota by leveraging counteracting inter-bacterial interactions, as we observed with the knockdown of *E. faecalis* and *B. fragilis*. These results suggest that phages with the appropriate properties could be leveraged as powerful tools to investigate the dynamics and interaction structure of the microbiome, providing either sustained knockdowns or precise transient perturbations.

Fascinatingly, our results reveal that phage predation in the gut microbiota has potential impact on the host, as manifested by modulation of the gut metabolome. With microbial metabolites having a substantial role in mediating the interaction between bacteria and the host (Dorrestein et al., 2014), the link between phage and microbial metabolites provides an interesting therapeutic avenue. Other methods of bacterial modulation, such as antibiotics can have profound and unpredictable results on microbial metabolism as demonstrated by streptomycin (Antunes et al., 2011) and cefoperazone (Theriot et al., 2014), which affected 87% and 53% of detected mouse fecal metabolites, respectively. By contrast, phage could elicit species-targeted effects as demonstrated with phage predation of *C. sporogenes* reducing tryptamine, which has been found to accelerate gastric mobility (Bhattarai et al., 2018), while predation of *E. faecalis* reduced tyramine, which can induce ileal contractions (Marcobal et al., 2012), for instance. Although much of microbial metabolism remains to be characterized (Dorrestein et al., 2014), the deployment of phage-directed bacterial knockdown may be a potential avenue for rationally modulating microbial metabolism for therapeutic purposes.

The longitudinal characterization of both phage and susceptible bacteria allowed us a view into the kinetics of phage predation in the gut microbiota. Among our findings is that lytic phage persists in the gut, the targeted bacteria experience knockdown but not eradication, and the proportion of phage to bacteria reveals dynamical insights within the gut. It has been proposed that the inability of phage to completely eradicate targeted gut bacteria is due to either genetic or ecological resistance mechanisms. Work has shown that genetic resistance may coincide with impaired fitness (Seed et al., 2014), allowing the susceptible strain to persist and thus propagate phage. Consequently, this fitness cost may explain why little evidence of co-evolution between bacterial resistance and phage predation was observed in the human gut virome (Reyes et al., 2010). Ecological resistance, where bacteria are physically inaccessible to phage, has also been proposed to explain why phage-targeted bacteria persist in the gut (Chibani-Chennoufi et al., 2004; Weiss et al., 2009; Zahid et al., 2008). While additional mechanistic studies would be valuable, the contrast in dynamics between the ratio of B40-8—*B. fragilis* (*i.e.*, an early amplification that peaks and decreases to steady state) and T4—*E. coli* (*i.e*., steady state) soon after phage administration may be indicative of their unique resistances to phage infection. For example, *B. fragilis* would be initially less accessible to phage as it colonizes the intestinal mucosa and whereas *E. coli* is more easily encountered by phage in the lumen (Lee et al., 2013).

The complexity and diversity of microbes and their interactions in the gut presents a tremendous experimental challenge. The conventional human gut microbiome is unique to each individual, composed of microbes that are often unculturable and difficult to phylogenetically classify, regularly perturbed by lifestyle, medication, dietary and environmental factors, and reciprocally influenced by the host. Therefore, to gain mechanistic insights into complex biological processes associated with the human gut microbiome, a compromise must be struck between how closely the model recapitulates the human gut and experimental pragmatism. Our use of a genetically inbred gnotobiotic mouse model colonized with ten culturable and characterized human commensal strains includes moderate complexity and bacterial diversity while minimizing potentially confounding variables. Our finding that lytic phages can play an unexpectedly broad and substantial role in the gut microbiota is only a glimmer of their potential functional impact as they have a diversity of lifestyles (*e.g.*, temperate), bacterial host ranges, and potential for horizontal gene transfer especially with regards to pathogenicity. By illuminating an important dynamical relationship between phage and commensal bacterial within the gut, we believe our work provides a framework that can guide future investigations of causal mechanisms between the microbiome and disease as well as the identification and development of future therapeutics.

## Acknowledgements

We are indebted to Richard Lavin, Nicholas DiBenedetto, Mary Delaney, and Andrea DuBois for training and advice in microbiological culture, and assistance with animal husbandry and anaerobic culture. We are grateful to Eli Bogart for insightful conversations. This project was funded by the Bill & Melinda Gates Foundation through the Grand Challenges Explorations Initiative (OPP1150555) and the Defense Advanced Research Program Agency (DARPA BRICS HR0011-15-C-0094).

## Author contributions

Conceptualization, B.B.H., T.E.G., and G.K.G.; Investigation, B.B.H., V.Y., and Q.L.; Resources, L.B., P.A.S., and G.K.G.; Writing – Original Draft, B.B.H.; Writing – Review & Editing, G.K.G. and P.A.S.; Funding Acquisition – B.B.H., P.A.S., G.K.G.

## Materials and Methods

### CONTACT FOR REAGENT AND RESOURCE SHARING

Further information and requests for resources and reagents should be directed to and will be fulfilled by the Lead Contact, Georg Gerber.

### EXPERIMENTAL MODEL AND SUBJECT DETAILS

#### Animal procedures

##### Ethical statement on mouse studies

All animal care and procedures were performed in accordance with institutional guidelines and approved by the Brigham and Women’s Hospital Institutional Animal Care and Use Committee under Protocol# 2017N000010.

##### Housing and husbandry of experimental animals

Germfree C57BL/6 mice were bred and maintained in isolators in the Massachusetts Host-Microbiome Center at Brigham and Women’s Hospital. Mice were provided water and double-irradiated standard chow *ad libitum* and maintained on a 12 hr light dark cycle.

For longitudinal phage perturbation experiments, five ∼8-10 week-old, male mice were individually-housed in a single sterile isolator and allowed to acclimated for 2 to 3 days. On experimental Day 0, mice were inoculated with a fresh preparation of 200 µL of defined bacterial consortia. This inoculate was prepared immediately prior to administration by thawing bacterial aliquots (previously snap-frozen and stored at −80°C), mixing at appropriate volumes for a final concentration of 10^8^ cfu/mL (*B. fragilis, B. ovatus, B. vulgatus, C. sporogenes, E. faecalis, E. coli, K. oxytoca*, and *P. distasonis*) or 10^7^ cfu/mL (*A. muciniphila* and *P. mirabilis*) of each species, and supplementing with BHI media. Samples were stored at −80°C. For phage administration on Days 16.08 and 30.08, mice received 100 µL of 1M sodium bicarbonate *per os* to neutralize gastric acid that was followed by 200 µL of phage mixture, 5 min later. Phages were combined immediately prior to administration from stock solutions stored in phage buffer at 4°C. On Day 16.1, each mouse received T4 and F1 phages (targeting *E. coli* and *C. sporogenes*, respectively) and on Day 30.1 each mouse received VD13 and B40-8 phages (targeting *E. faecalis* and *B. fragilis*, respectively). Additional phage buffer was supplemented for a final concentration of 10^7^ pfu/mL of each phage. Stool samples were collected by placing each mouse into a sterile beaker, collecting the freshly voided feces, and snap-freezing in liquid nitrogen within 30 min. Stool for each mouse was collected at these time points: 0.26, 0.48, 1.08, 1.31, 2.08, 2.31, 3.06, 5.13, 7.06, 8.06, 9.04, 11.04, 13.27, 14.08, 15.08, 16.06, 16.31, 17.06, 17.27, 18.06, 18.27, 13.08, 21.07, 23.08, 25.08, 27.06, 28.08, 29.08, 30.38, 30.52, 31.08, 31.25, 32.08, 32.31, 33.29, 35.10, 37.06, 39.06, 40.15, 42.10, 43.06 days.

Dropout experiments were conducted similarly as above with specific differences. Twenty ∼8-10 week-old, male mice were individually-housed and randomly assigned into four groups with each group in separate sterile isolators. On experimental Day 0, each set of mice were inoculated with 200 µL of a dropout mixture consisting of the defined bacterial consortia without one of the following species: *B. fragilis, C. sporogenes, E. faecalis*, or *E. coli.* Stool was collected by the previously described method at these time points: 0.26, 1.1, 3.1, 7.1, 11.1, and 16.1 days.

##### Bacterial strains

Bacteria were cultured from single colonies at 37°C either in BHI under aerobic conditions (*E. coli, E. faecalis, K. oxytoca*, and *P. mirabilis*) or in BHI (+vitamin K, +hemin) under anaerobic conditions (*A. muciniphila, C. sporogenes B. fragilis, B. ovatus, B. vulgatus*, and *P. distasonis*). After incubation, *B. fragilis*, *C. sporogenes*, and *E. coli* were concentrated ∼8- to 10-fold by centrifugation. All cultures were snap-frozen in liquid nitrogen and stored at −80°C. Each batch was checked for titer and contamination by diluting aliquots in PBS and culturing on non-selective plates (TSA with 5% sheep blood for aerobic culture or Brucella agar with 5% sheep blood, hemin and vitamin K for anaerobic culture). *P. mirabilis* dilutions were also cultured on MacConkey agar plates to inhibit swarming so that individual colonies could be counted.

##### Phage strains

High titer phage stocks were propagated on their host bacteria using a soft-agar overlay technique by mixing 100 µL dilutions of phage in phage buffer with 100 µL of an overnight bacterial culture, adding 3 mL of molten nutrient soft-agar at ∼42°C and 30 µL of 1 M calcium chloride, then immediately pouring onto petri dishes of the same nutrient agar and allowed to harden at room temperature. Nutrient agar (1.5% agar) and soft agar (0.3% agar) consisted of BHI (for VD13 phage on *E. faecalis*), TNT (for T4 phage on *E. coli*), or BHI +vitamin K +hemin (for F1 phage on *C. sporogenes* and B40-8 phage on *B. fragilis*). Plates were incubated overnight at 37°C in air for phage propagated on facultative anaerobes (*E. faecalis* and *E. coli*) or anaerobically for phage propagated on obligate anaerobes (*B. fragilis* and *C. sporogenes*). Phage was harvested from plates showing the greatest density of plaques by resuspending the soft-agar overlays into 10 mL of phage buffer, gently rocking at 4°C for ∼2 h to extract phage and then centrifuging at 4000 rpm at 4°C and sterile filtering the supernatant. Phage titers were measured via plaque assay using the same soft-agar overlay technique.

Phage infectivity (i.e., bacterial host range) was determined by a spot titer technique (Kutter, 2009) where 5 µL spots of ∼10^9^ pfu/mL of each phage (or phage buffer as a control) was added to soft-agar overlays of each bacteria (without phage), prepared as described above. Plates were incubated overnight at 37°C aerobically or anaerobically depending on the bacterial culture conditions.

##### Nucleic acid extraction

Extraction of DNA from fecal samples was performed using the ZymoResearch ZymoBIOMICS DNA 96-well kit according to manufacturer instructions.

##### 16s rRNA amplicon sequencing

PCR amplification of the V4 region of the 16s rRNA gene was conducted following a previously described protocol using universal bacterial primers (515F and 806R) with dual-index barcodes (Kozich et al., 2013):

> 5’-[Illumina adaptor]-[unique barcode]-[sequencing primer pad]-[linker]-[primer]

Read 1 (fwd primer): AATGATACGGCGACCACCGAGATCTACAC-NNNNNNNN-TATGGTAATT-GT-GTGCCAGCMGCCGCGGTAA
Read 2 (rev primer): CAAGCAGAAGACGGCATACGAGAT-NNNNNNNN-AGTCAGTCAG-CC-GGACTACHVGGGTWTCTAAT

After successful amplification was determined by the presence of a 384 bp band on a 1.5% agarose gel, concentration was measured using a Quan-IT dsDNA high sensitivity assay (Invitrogen). Roughly 120 ng of each amplification product was pooled to generate an aggregated library, from which 300-500 bp amplicons were selected using a targeted size selection platform, Pippin Prep with 1.5% agarose cassette (Sage Sciences) according to the manufacturer’s instructions. Amplicon size was characterized with an Agilent Technologies 2100 bioanalyzer trace. The aggregated library was denatured with sodium hydroxide and diluted to 7.5 pM in HT buffer (Illumina). 480 µL was then combined with 120 µL of 7.5 pM phiX and loaded onto a MiSeq V2 reagent cartridge (Illumina). Sequencing was conducted with 250 bp paired-end reads using custom sequencing primers:

> 5’-[sequencing primer pad]-[linker]-[primer]

Read 1: TATGGTAATT-GT-GTGCCAGCMGCCGCGGTAA
Read 2: AGTCAGTCAG-CC-GGACTACHVGGGTWTCTAAT

> 5’-[primer]-[linker]-[sequencing primer pad]

Index primer: ATTAGAWACCCBDGTAGTCC-GG-CTGACTGACT

MiSeq sequencing was performed using the default parameters and standard operating procedures for Illumina MiSeq operation to generate a demultiplexed fastq files. The raw fastq files containing sequencing reads were subjected to a quality control procedure described below. fastqc(v0.11.2) (Andrews) was used to check read quality. Trimmomatic (v0.36) (Bolger et al., 2014) was used to trim low quality bases and filter very short reads (<120 nt) after trimming. Read depth of time-series data was 63,435 ± 16,343 and for bacterial knockout data was 66,683 ± 19,204. Calculation of relative abundances accounted for species-specific number of 16s rRNA copies per genome (Stoddard et al., 2015).

##### qPCR protocol for each phage and total bacteria

Each phage was quantified with phage-specific primers (10 µM) using the PowerUP SYBR Green Master Mix (ThermoFisher) according to manufacturer instructions. Template DNA from fecal extracts and liquid culture standards (quantified by plaque assay) were diluted 100-fold for measurement.

Total bacteria was quantified from extracted DNA using a TaqMan Universal Master Mix II no UNG kit (ThermoFisher 4440040) with the TaqMan Gene Expression Assay (ThermoFisher 4331182) primer and probe (FAM-MGB) set for 16s rRNA quantification (ThermoFisher assay ID Pa04230899_s1) and performed according to manufacturer instructions. Liquid cultures of *E. coli* Nissle 1917 (quantified by plating and colony counting) were used as standards.

Measurements were performed in 96-well or 384-well plates using an Applied Biosystems QuantStudio 12k Flex Real-Time PCR system. For instances where certain bacterial species were transiently undetectable in the time-series or knockout animal experiments, values were set to the lowest otherwise detected concentration in our data set: [*C. sporogenes*]_min_ = 1.58 cfu/g stool; [*E. faecalis*]_min_ = 5.3 × 10^2^ cfu/g stool; [*P. distasonis*]_min_ = 5.5 cfu/g stool.

##### Metabolomics

Fecal samples obtained as described above and stored at −80°C were delivered to Metabolon (Durham, NC USA) where sample preparation and analysis was performed. Samples (∼1-2 pellets) were homogenized in methanol at 50 mg/mL for metabolite extraction. The supernatant was separated from debris and precipitates (*e.g.*, proteins) by centrifugation, divided into five aliquots for four different analysis conditions plus one backup sample, and placed into a TurboVap (Zymark) for solvent removal. Dried samples were stored under nitrogen gas overnight until analysis.

All samples were reconstituted and measured using a Waters ACQUITY ultra-performance liquid chromatography (UPLC) instrument with attached Thermo Scientific Q-Exactive high resolution/accurate mass spectrometry (MS), heated electrospray ionization source (HESI-II), and Orbitrap mass analyzer (35,000 mass resolution), as similarly described previously (Am et al., 2014). Each of four aliquots were analyzed as follows: (1) elution with a C18 column (Waters UPLC BEH C18-2.1×100mm, 1.7µm) in positive-ion mode with a water/methanol gradient mobile phase containing 0.05% perfluorpentanoic acid (PFPA) and 0.1% formic acid (FA); (2) similarly to the previous method except with a water/acetonitrile/methanol gradient mobile phase containing 0.05% PFPA and 0.01% FA; (3) elution with a separate C18 column in negative-ion mode with a water/methanol gradient mobile phase containing 6.5 mM ammonium bicarbonate, pH 8; (4) elution with a HILIC column (Waters UPLC BEH amide 2.1×150mm, 1.7µm) in negative-ion mode with a water/acetonitritile gradient mobile phase containing 10 mM ammonium formate, pH 10.8. MS analysis utilized dynamic exclusion, alternating between MS and data-dependent MS^n^ scans. Scan range covered 70-1000 m/z. Data extraction, peak-identification, and quality control was conducted using Metabolon’s proprietary software. Compounds were identified and quantified by comparison to a library of standards.

